# Neuronal activity-related transcription is blunted in immature compared to mature dentate granule cells

**DOI:** 10.1101/2022.09.26.508896

**Authors:** Sarah L Parylak, Fan Qiu, Sara B Linker, Iryna S Gallina, Christina K Lim, David Preciado, Aidan H McDonald, Fred H Gage

## Abstract

Immature dentate granule cells (DGCs) generated in the hippocampus during adulthood are believed to play a unique role in dentate gyrus function. Although immature DGCs have hyperexcitable membrane properties in vitro, the consequences of this hyperexcitability in vivo remain unclear. In particular, the relationship between experiences that activate the dentate gyrus, such as exploration of a novel environment (NE), and downstream molecular processes that modify dentate gyrus circuitry in response to cellular activation are unknown in this cell population. We first performed quantification of immediate early gene proteins in immature (5-week-old) and mature (13-week-old) DGCs from mice exposed to a NE. Paradoxically, we observed lower immediate early gene protein expression in hyperexcitable immature DGCs. We then isolated nuclei from active and inactive immature DGCs and performed single nuclei RNA-Sequencing. Compared to mature nuclei collected from the same animal, immature DGC nuclei showed less activity-induced transcriptional change, even though they were classified as active based on expression of ARC protein. These results demonstrate that the coupling of spatial exploration, cellular activation, and transcriptional change differs between immature and mature DGCs, with blunted activity-induced changes in immature cells.

## Introduction

The dentate gyrus (DG) of the mammalian hippocampus continues to produce new neurons even in adulthood (Kempermann et al., 2015). The precise contribution of adult-born neurons to the function of the hippocampus remains a subject of debate and may differ depending on these neurons’ developmental stage. Within a few months of their birth in the rodent hippocampus, adult-born dentate granule cells (DGCs) reach maturity and attain morphological and physiological features that are generally considered indistinguishable from their developmentally generated counterparts (Laplagne et al., 2006; van Praag et al., 2002). During their maturation process, however, immature DGCs possess distinct electrophysiological properties.

Immature DGCs under 6-7 weeks of age show an intrinsic electrophysiological hyperexcitability. They are characterized by high input resistance, high efficacy of firing to current injection, and increased activation in response to perforant path stimulation (Dieni et al., 2016; Marín-Burgin et al., 2012; Mongiat et al., 2009; Yang et al., 2015). These immature cells also show enhanced long-term potentiation (LTP), with a reduced threshold for induction and increased amplitude (Ge et al., 2007; Schmidt-Hieber et al., 2004). Reduced synaptic connectivity may counterbalance cellular hyperexcitability, at least in part. Immature DGCs are more limited by excitatory drive than mature cells, and stimulation of the entorhinal cortex has been reported to poorly recruit immature DGCs (Dieni et al., 2013, 2016). While in vivo data on the firing patterns of immature cells are limited, immature DGCs were shown to generate more calcium transients than mature cells in one study using head-fixed imaging (Danielson et al., 2016). Combined, these results have led to considerable speculation about how immature DGCs contribute to the function of the DG in learning and memory.

Memory formation requires not just synaptic activity, but also new transcription and translation (Alberini & Kandel, 2015; Costa-Mattioli et al., 2009; Yap & Greenberg, 2018). An initial wave of activity-induced transcription is detectable within minutes of a stimulus and results in the expression of immediate early genes (IEGs) such as *Fos* and *Arc* (Yap & Greenberg, 2018). Additional transcriptional changes occur over the next several hours, and transcriptional and translational inhibitors can continue to impair memory over 1-2 days (Alberini & Kandel, 2015; Costa-Mattioli et al., 2009; Yap & Greenberg, 2018). To explore the transcriptional mechanisms downstream of neuronal activity in the DG in an unbiased manner, we previously used single nuclei RNA sequencing (snRNA-Seq). We exposed mice to a NE, isolated individual DGCs, and classified them as active or inactive based on the expression of IEG proteins (Jaeger et al., 2018; Lacar et al., 2016). We observed that DGCs, whose firing and IEG expression are quite sparse in comparison to other hippocampal cell types, have a more dramatic transcriptional response to activation than CA1 pyramidal cells (Jaeger et al., 2018). Enhanced activity-dependent gene expression in DGCs may support the development of highly selective responses to specific stimuli. However, our previous work did not address the contribution of immature cells.

In the current study, we sought to compare activity-related changes in immature and mature DGCs by addressing 2 questions. First, are immature DGCs more or less likely than mature DGCs to express IEG markers of activation in response to a NE? Second, among immature and mature DGCs that express IEG proteins, what transcriptional changes are observed in active cells? We hypothesized that hyperexcitable immature DGCs would show enhanced IEG expression and activity-dependent transcriptional change. Counterintuitively, we observed that immature DGCs are both less likely to show IEG markers of activity and less transcriptionally modified after activation.

## Results

We previously studied how neuronal activation arising from NE exposure (Jaeger et al., 2018; Lacar et al., 2016) triggered activity-related transcription in DGCs, but we did not discriminate between immature and mature DGCs. To identify immature and mature DGCs, cells were birthdated in adult mice expressing a nuclear membrane-targeted GFP under the control of an Ascl1-driven Cre recombinase (Ascl1CreERT2 x LSL-Sun1-sfGFP mice, Fig 1A). In response to tamoxifen (TAM) treatment, GFP is expressed in Ascl1+ neural progenitors within the DG, and the GFP signal persists in cells as they differentiate and mature, remaining clearly visible within the subgranular zone and granule cell layer up to 13 weeks later (Fig 1B).

**Figure 1.**
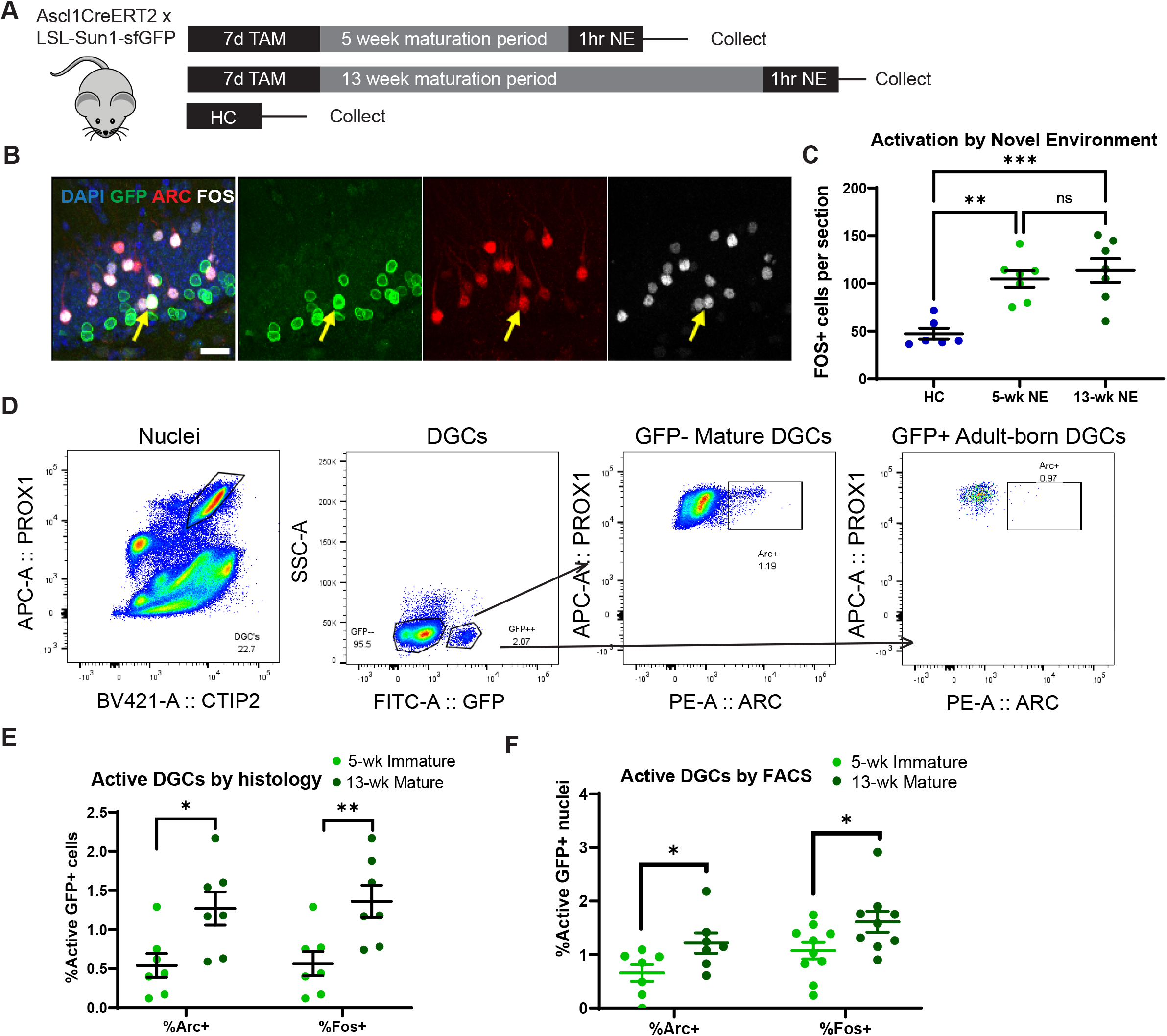
Reduced immediate early gene (IEG) protein expression in immature DGCs. A) Experimental timeline. Mice received tamoxifen (TAM) gavage for 7 days to label DGCs and were exposed to a novel environment (NE) to stimulate activation 5 or 13 weeks (wk) later. Untreated littermates collected directly from the home cage (HC) were used as a control. B) Sample of durable GFP labeling 13 weeks after TAM. Scale bar 20 um. Yellow arrow indicates a cell triple labeled for GFP, ARC, and FOS. C) NE exposure induces significantly more FOS and ARC expression than the home cage control condition (n’s: HC = 6, 5-wk NE = 7, 13-wk NE = 7). D) Sample flow cytometry (FACS) gating strategy for identifying active immature DGCs. DGCs were identified by co-labeling with marker proteins PROX1 and CTIP2. Cell age was identified by GFP. Cell activation was assessed with ARC or FOS. E) Quantification of active 5-week and 13-week DGCs in coronal tissue sections shows greater IEG expression in more mature cells (n’s: 5-wk = 7, 13-wk = 7). F) Quantification of active 5-week and 13-week DGCs via FACS also shows greater IEG expression in more mature cells (n’s: Arc 5-wk = 7, Arc 13-wk = 7, Fos 5-wk = 10, Fos 13-wk = 9). Error bars are mean +/- S.E.M. *p<0.05, **p<0.01, ***p<0.001, ns = not significant.

Because immature DGCs are electrophysiologically hyperexcitable, the ability of this immature population to express IEGs in response to a behaviorally relevant stimulation was measured and compared to mature cells. Ascl1CreERT2 x LSL-Sun1-sfGFP mice were treated with TAM to label newborn cells and exposed to a NE 5 or 13 weeks later, when labeled cells were immature or mature, respectively (Fig 1A). Mice were perfused immediately after NE exposure, and expression of IEG proteins was quantified in coronal sections of the DG. Untreated wildtype littermates collected directly from their home cages were used as a control. As we previously observed, 1 hour of NE exposure led to a significant increase in FOS+ cells compared to home cage controls (HC) (Fig 1C; one-way ANOVA F(2,17)=13.24, p=0.0003, Tukey’s multiple comparisons test p=0.0019 for HC vs. 5-week, p=0.0005 for HC vs. 13-week, p=ns for 5-week vs. 13-week). Surprisingly, both of these IEG proteins were expressed at a lower rate in immature (5 week) compared to mature (13 week) GFP+ cells (Fig 1E; ARC: Welch’s t-test t=2.797, p=0.0176; FOS: Welch’s t-test t=3.103, p=0.0099). Activation rates of immature and mature GFP+ cells were also compared using flow cytometry (Fig 1D, 1F). At 5 weeks of age, GFP+ nuclei expressed ARC and FOS at a reduced rate compared to mature 13-week-old nuclei (Fig 1F; ARC: Welch’s t-test t=2.261, p=0.0438; FOS: Welch’s t-test t=2.166, p=0.0461). These results indicate that immature DGCs, despite their electrophysiological hyperexcitability, are actually less likely to express protein markers of activation than mature DGCs. The vast majority of ARC+ and FOS+ cells observed in the DG during memory encoding are thus mature DGCs.

We next asked whether different activity-related transcriptional programs were triggered in mature DGCs and the subset of immature DGCs that did successfully generate ARC protein in response to a NE. As above, newborn cells were labeled in Ascl1CreERT2 x LSL-Sun1-sfGFP mice and allowed to mature to 5 weeks of age. Mice were exposed to a NE for 1 hour, then sacrificed for hippocampus dissection either immediately (1-hr group) or after 3 additional hours in the home cage (4-hr group) (Fig 2A). These time points were selected to capture a first wave of transcription corresponding to IEG expression and a second wave corresponding to late response genes. The dissected hippocampus was dounce homogenized and stained for DGC marker proteins PROX1 and CTIP2, cell age marker GFP, and activation marker ARC. Nuclei from 4 populations were isolated for sequencing (Fig 2B) at each of these timepoints: 1) Mature Inactive cells; 2) Immature Inactive cells; 3) Mature Active cells; and 4) Immature Active cells.

**Figure 2.**
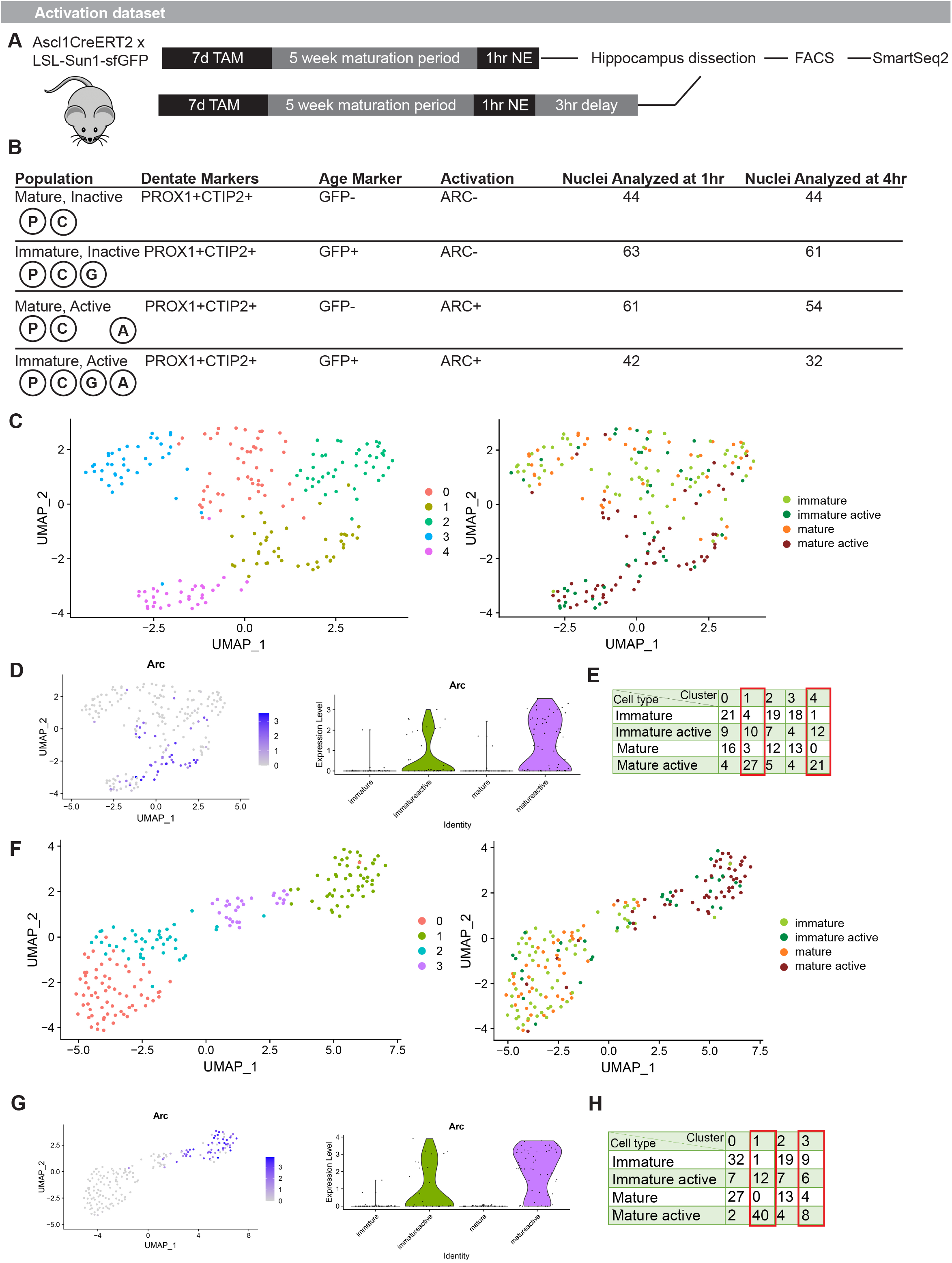
Reduced transcriptional response to activation in immature DG neurons. A) Experimental timeline. Mice received TAM to label DGCs, were exposed to a NE 5 weeks later, and then were sacrificed for hippocampus dissection and nuclei preparation immediately or after an additional 3-hour delay. B) Diagram of nuclear cell markers used to isolate specific populations via FACS and total cells per population included in the analysis. C) Unbiased clustering of nuclei isolated at 1 hour after the start of NE. D) *Arc* expression in nuclei isolated at 1 hour identifies active clusters. E) Mature Active cells preferentially localize to clusters 1 and 4. Immature Active cells are more broadly distributed across clusters despite isolation based on ARC protein expression. F) Unbiased clustering of nuclei isolated at 4 hours after the start of NE. G) *Arc* expression in nuclei isolated at 4 hours identifies active clusters. H) Mature Active cells preferentially localize to clusters 1 and 3. Immature Active cells are more broadly distributed across clusters despite isolation based on ARC protein expression.

As suggested by the above histology and flow cytometry quantifications in 5-vs. 13-week cells, the Immature Active population was extremely small (Fig S1). After nuclei isolation and RNA-Seq library preparation by SmartSeq2, these 4 populations were compared within each time point. Prior work in our lab has demonstrated that the activity-related transcriptional response differs at 1 and 4 hours after the activation event, as some IEGs return to baseline levels, others exhibit sustained changes at both time points, and some late response genes are activated only at the 4-hour time point.

At 1 hour after activation, unbiased clustering segregated most Mature Active cells, featuring ARC protein expression, into 2 clusters that also exhibited high relative levels of *Arc* transcripts (Fig 2C-D). As expected, few Mature Inactive or Immature Inactive cells were found within these active clusters. In contrast, Immature Active cells did not preferentially localize with Mature Active cells. Instead, Immature Active cells were broadly distributed across clusters (Fig 2E; Fisher’s exact test for Mature Active vs. Immature Active in active clusters 1, 4 vs. inactive clusters 0, 2, 3, p=0.0093). Similar results were observed at 4 hours after activation. Again, the vast majority of Mature Active cells identified by ARC protein expression occupied active clusters with higher levels of *Arc* transcripts (Fig 2F-G). Immature Active cells were again broadly distributed across clusters instead of preferentially belonging to active clusters (Fig 2H; Fisher’s exact test for Mature Active vs. Immature Active in active clusters 1, 3 vs. inactive clusters 0, 2, p=0.0011). These results suggest that, even among the 5-week-old DGCs that activated in response to a NE, the overall transcriptional response to activation was blunted relative to the response observed in mature cells.

Further analysis of active vs. inactive cells revealed that the number of differentially expressed genes detected was far greater in mature than immature cells at both the 1- and 4-hour time points (Fig 3A). Among the differentially expressed genes, 3 were common to both immature and mature cells at 1 hour and 5 were common to both immature and mature cells at 4 hours (Fig 3B). The remaining differentially expressed genes were unique to mature cells. No differentially expressed genes were unique to the Immature Active vs. Immature Inactive comparison. Activation in mature cells significantly upregulated the expression of IEGs previously identified in active DGCs such as *Homer1* and *Bdnf* by 1 hour, but these genes fell short of significance in immature cells (Fig 3C). Sustained upregulation of some genes, including *Bdnf, Spry2*, and *Pim1*, was observed only in mature cells at 4 hours, along with late response genes we previously identified in active DGCs such as *Sorcs3* and *Blnk*. GO-term analysis of the activity-response genes unique to mature cells showed significant enrichment of terms related to postsynaptic membrane, cell junction, and postsynaptic density (Fig 3D-E). Overall, these results demonstrate that activity-related transcriptional change in the DG occurs mainly in mature cells. Among immature DGCs sufficiently active to express ARC protein, their transcriptional state is activated to a lesser degree than mature cells, failing to show the same level of activity-induced gene expression. These findings suggest that mature cells are the main substrate for cellular modification in response to memory encoding, whereas immature cells contribute via indirect effects on DG network activity (Fig 3F).

**Figure 3.**
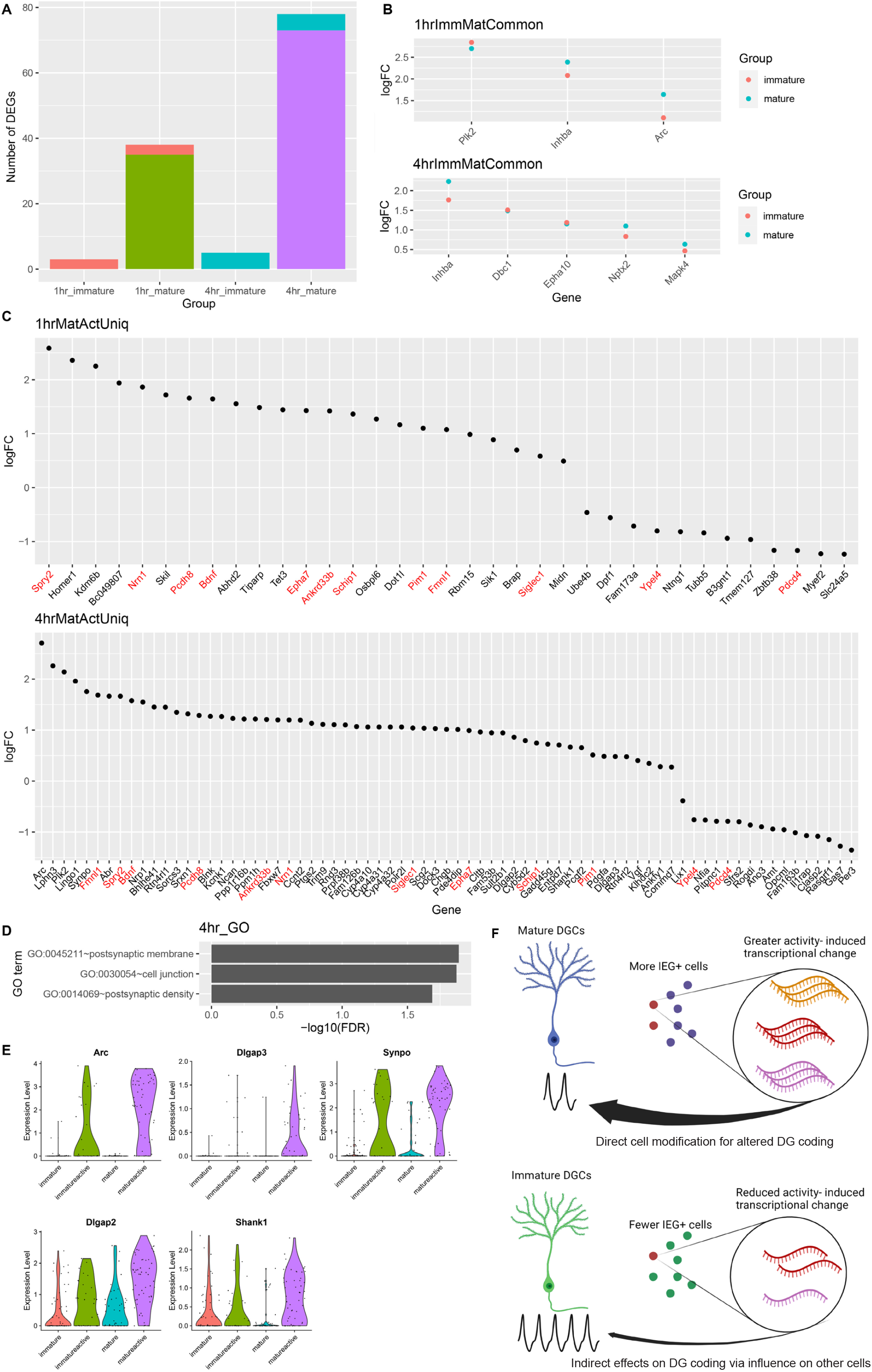
Immature DG neurons show activity-related changes in transcription of only a subset of genes observed in mature. A) Number of differentially expressed genes (DEGs) between Active and Inactive nuclei based on maturation status and time point, e.g. 1-hour immature indicates the number of DEGs between Immature Active and Immature Inactive cells isolated at 1 hour after start of NE. Coral bar indicates genes shared between 1-hour immature and 1-hour mature comparisons. Blue bar indicates genes shared between 4-hour-immature and 4-hour mature comparisons. B) Log2 fold change (logFC) between active and inactive nuclei of DEGs common to both immature and mature nuclei from A. C) Fold change of DEGs uniquely identified in mature, but not immature, active and inactive nuclei comparisons. DEGs detected in mature cells at both 1 and 4 hour after the start of NE are labeled in red. No DEGs were uniquely identified in immature cells. D) GO terms significantly enriched among DEGs uniquely identified in mature cells 4 hours after activation. E) Violin plots of DEGs presented in both “postsynaptic membrane” and “postsynaptic density” terms from D. F) Model for contribution of immature DGCs to activity-dependent changes (created with BioRender.com). Mature cells respond by engaging cell intrinsic mechanisms of transcriptional change, whereas physiologically hyperactive immature cells may contribute more indirectly by influencing other cells.

## Discussion

We observed reduced in vivo recruitment of immature 5-week-old DGCs compared to more mature cells when they were stimulated by NE exploration. Even among those 5-week-old DGCs that expressed activation marker ARC, activity-induced transcriptional change in immature DGCs was less pronounced than in their mature counterparts. These results were surprising, given the consistent finding from electrophysiological studies that immature DGCs under about 6-7 weeks of age are hyperexcitable to perforant path stimulation or direct current injection in ex vivo slices. Functionally, our results demonstrate that the coupling of behaviorally induced cortical input to the DG, expression of IEG proteins, and downstream consequences of IEG expression differ between mature and immature cells. These differences further suggest that the role of immature DGCs is likely to be an indirect one. Immature DGCs may contribute to hippocampal function by modifying the response of mature cells rather than in undergoing dramatic activity-dependent changes to directly encode a memory engram.

We decided to focus on 5-week-old DGCs to target an age when immature neurons are synaptically connected both via inputs from the entorhinal cortex and outputs to CA3, but still show unique physiological properties (Dieni et al., 2016; Ge et al., 2007; Toni et al., 2008; Yang et al., 2015; Zhao et al., 2006). We thus could have missed an effect that occurs earlier in their development. However, given previous studies showing minimal IEG expression prior to 3-4 weeks of age (Jessberger & Kempermann, 2003; Jungenitz et al., 2014; Snyder et al., 2009), this effect would have to be quite transient. A few papers have reported increased in vivo recruitment of immature DGCs at age ranges similar to those studied here (Kee et al., 2007), but other groups have published conflicting results (Snyder et al., 2009; Stone et al., 2011). Our results add to literature suggesting that immature DGCs are gradually incorporated into the circuitry of the DG, with synaptic connectivity and IEG expression converging on mature levels over a period of 1-2 months, rather than showing a hyperactive phase of IEG induction.

This study is, to the best of our knowledge, the first to use snRNA-Seq to compare activity-dependent transcription in adult-born DGCs. We elected to study nuclei to avoid inconsistent recovery of dendritic compartments and to prevent spurious IEG expression from whole-cell dissociation methods (Lacar et al., 2016; Wu et al., 2017). However, our approach will miss differences in dendritic RNA. It is possible that immature and mature DGCs differ in RNA localization either at rest or following activation. Additionally, the low fraction of immature DGCs within the dentate compared to mature DGCs, combined with their lower rate of ARC and FOS expression, means that IEG-positive immature DGCs are extraordinarily rare. The rarity of this population makes it challenging to obtain large sample sizes or use droplet-based sequencing methods to tease out any subpopulations of immature DGCs with different levels of responsiveness. Other stimulation methods, such as seizure-inducing drugs, optogenetics, or chemogenetics, may be valuable in future studies to obtain larger populations for analysis. However, these methods all have the tradeoff of being less reflective of endogenous activation processes.

The functional role of immature DGCs remains a subject of active debate. Although our results show that activity-dependent transcription in immature DGCs is reduced, prior studies demonstrate that immature DGCs still make an important functional contribution. Immature DGCs influence the ability to discriminate between similar experiences, known as behavioral pattern separation, as well as mood-related behaviors (Anacker & Hen, 2017; Toda et al., 2019). Exactly how this influence occurs is unclear. It has been hypothesized that the role of hyperexcitable immature DGCs is not to carry a specific signal but rather to modulate the activity of mature cells in the DG (Anacker & Hen, 2017; Drew et al., 2016). Several groups have reported that a loss of immature DGCs increases activation of the DG, including greater spread of current injection via the perforant path or greater IEG activation (Anacker et al., 2018; Burghardt et al., 2012; Ikrar et al., 2013). If this modulatory effect occurs at levels of activity that are insufficient to trigger IEG expression, it may explain how immature DGCs can be physiologically hyperactive without showing the same downstream modifications observed in mature DGCs. Even DGCs within the first 2-3 weeks of life can be stimulated to produce spikes, prior to any reliable induction of IEG proteins (Li et al., 2017; Schmidt-Hieber et al., 2004). Action potentials may thus still be generated in immature DGCs, allowing them to recruit inhibitory feedback via hilar interneurons, but these action potentials may not be converted into IEG expression as efficiently as in mature cells.

Immature DGCs may also provide a response that is important due to its quality rather than its quantity. Immature DGCs, whose developing dendritic arbors are smaller than those of mature cells, may provide less correlated output (Dieni et al., 2016; Li et al., 2017). Less correlated output would in turn provide unique computational power to the DG circuit even if the transcriptional response is reduced. Immature DGCs may also have a longer-term impact on DG circuitry as they compete for synaptic connections with existing cells (Akers et al., 2014; McAvoy et al., 2016). Environmental conditions as immature DGCs are developing are known to influence their survival (Tashiro et al., 2007), and different cohorts of immature DGCs maturing over different intervals may add temporal information not detectable over the short time course examined here (Rangel et al., 2014). Further studies of activity-dependent transcription under conditions where neurogenesis is ablated or enhanced could provide evidence of how this small fraction of cells can influence the DG and related behaviors.

## Methods

### Animals and drug treatment

All animal procedures were approved by the Institutional Animal Care and Use Committee of The Salk Institute for Biological Studies. Experiments were conducted in accordance with the National Institutes of Health’s Guide for the Care and Use of Laboratory Animals and the US Public Health Service’s Policy on Humane Care and Use of Laboratory Animals. To generate Ascl1CreERT2 x LSL-Sun1-sfGFP mice, hemizygous Ascl1^tm1.1(Cre/ERT2)Jejo^/J mice (Jackson Laboratory stock #012882, RRID: IMSR_JAX:012882) (Kim et al., 2011) were bred with homozygous B6;129-Gt(ROSA)26Sor^tm5(CAG-Sun1/sfGFP)Nat^/MmbeJ (Jackson laboratory stock #030952, RRID:IMSR_JAX:030952, a generous gift from Dr. Margarita Behrens) (Mo et al., 2015). All mice were housed under a standard 12-hr light/dark cycle with *ad libitum* access to rodent chow and water. Male and female mice were 6-10 weeks of age at the beginning of drug treatment. To induce Cre-recombination and label immature cells, tamoxifen (Sigma, T5648) was delivered at 200 mg/kg in 90% corn oil/10% ethanol by oral gavage daily for 7 days.

### NE exposure

To stimulate behaviorally relevant neuronal activity, mice were exposed to a NE for 1 hour. NE cages contained huts, tunnels, running wheels, and sawdust bedding or a textured plastic floor. Cage dimensions were 54″ x 34″ base, 12″ height or 42″ x 42″ base, 12″ height. At the conclusion of NE exposure, mice were returned to the home cage until sacrifice.

### Histology

For histological analysis, mice were deeply anesthetized with ketamine/xylazine/acepromazine i.p. and transcardially perfused with 0.9 % NaCl followed by 4% paraformaldehyde (PFA). Brains were extracted, post-fixed in PFA overnight, and then sunk in 30% sucrose. Coronal sections (20-40 um) were obtained on a sliding microtome and stored at -20C until staining. For staining, a 1 in 12 series of sections were washed in TBS, blocked with 0.25% Triton X-100 in TBS with 3% horse serum and incubated with primary antibody diluted in blocking buffer for 3 nights at 4C. Sections were washed and incubated with fluorophore-conjugated secondary antibodies for 2 hours at room temperature. DAPI was applied at 1:1000 in TBS wash following secondary incubation. At the conclusion of TBS washes, sections were mounted with Immu-Mount mounting media.

### Antibodies

#### Antibodies used in this study were

Mouse anti-PROX1 (EMD Millipore #MAB5654, RRID:AB_2170714)

Rat anti-CTIP2 (Abcam #ab18465, RRID:AB_2064130)

Chicken anti-GFP (Aves Labs #GFP-1020, RRID:AB_10000240)

Guinea pig anti-ARC (Synaptic Systems #156005, RRID:AB_2151848 and #156004, RRID:AB_2619853)

Rabbit anti-FOS (Synaptic Systems #226003, RRID:AB_2231974)

Donkey anti-mouse AF647 (Jackson Immuno Research #715-605-151, RRID:AB_2340863)

Donkey anti-rat Dyl405 (Jackson Immuno Research #712-475-153, RRID:AB_2340681)

Donkey anti-chicken AF488 (Jackson Immuno Research #703-545-155, RRID:AB_2340375)

Donkey anti-guinea pig Cy3 (Jackson Immuno Research #706-165-148, RRID:AB_2340460)

Donkey anti-rabbit Cy3 (Jackson Immuno Research #711-165-152, RRID:AB_2307443)

Donkey anti-rabbit AF647 (Jackson Immuno Research #711-605-152, RRID:AB_2492288)

### Imaging and quantification

Images of the DG were obtained on a Zeiss laser scanning confocal microscope (710 or 880) using a 20x objective. Z-stacks were obtained through the full section thickness, and tiles spanning the DG were stitched using ZEN software. Active cells (ARC+ or FOS+) and colocalization with markers of maturity (GFP) were quantified manually by a blinded observer.

### Nuclei dissociation

Nuclei dissociation was performed as previously described (Jaeger et al., 2018; Lacar et al., 2016). Briefly, the hippocampus was dissected bilaterally after deep anesthesia with ketamine/xylazine/acepromazine and cervical dislocation. Hippocampal tissue was immediately placed into a nuclei isolation medium (sucrose 0.25 M, KCl 25 mM, MgCl2 5 mM, TrisCl 10 mM, dithiothreitol 100 uM, 0.1% Triton, protease inhibitors). Tissue was dounce homogenized and then samples were washed, resuspended in nuclei storage buffer (0.167 M sucrose, MgCl2 5 mM, and TrisCl 10 mM, dithiothreitol 100 uM, protease inhibitors) and filtered. Antibody staining proceeded in the nuclei storage buffer. Solutions and samples were kept cold throughout the protocol. For RNA-seq experiments, tools and solutions were made RNAse-free and RNAse inhibitors were used (Ambion #AM2684 at 1:1000 in both isolation and storage buffers).

### Flow cytometry

For analysis of activation rates in immature and mature cells, dissociated nuclei were stained for dentate markers PROX1 and CTIP2, cell age indicator GFP, and the IEGs ARC or FOS. Data was collected on a BD LSRII cytometer (BD Biosciences) and analyzed with FlowJo software (Tree Star). Nuclei were gated based on forward and side scatter, and DGCs were identified based on co-labeling with PROX1 and CTIP2. Nuclei from mice lacking the Ascl1CreERT2 transgene and not exposed to the NE were used as negative controls to validate gating strategy for immature (GFP+) and active (ARC or FOS+) cells. For nuclei isolation prior to RNA-Seq, samples were run on a BD Influx cell sorter and collected directly into 384-well plates pre-filled with 1 uL lysis buffer (0.1% Triton-X, 1 U/ul RNAse inhibitor, 2.5 mM dNTP mix, 2.5 uM anchored oligo-dT primer) and frozen at -80C until further processing.

### Single-nuclei RNA-seq

Libraries were prepared using the Smart-seq2 protocol as previously described with minor modifications (Jaeger et al., 2018; Picelli et al., 2013, 2014). First strand synthesis with ProtoScript II reverse transcriptase (New England Biolabs), PCR preamplification with KAPA Hifi HotStart ReadyMix (Kapa), and bead cleanup (Agencourt AMPure XP beads, Beckman Coulter) were performed with the aid of a Mosquito HV liquid handler (SPT Labtech). The quality of cDNA was assessed by Bioanalyzer chip (Agilent) and concentration quantified by Qubit assay (Invitrogen). Single nuclei libraries were prepared with a Nextera XT DNA library prep kit (Illumina) and pooled together for sequencing. The quality of pooled libraries was assessed by TapeStation assay (Agilent). Libraries were sequenced on an Illumina HiSeq using single-end 50bp reads by the Salk Institute Next Generation Sequencing Core. Reads were trimmed using Solexa-QA++ dynamic trim and aligned to the mm10 (GRCm38) reference genome with Ensembl gene annotation using RSEM (bowtie). Transcripts per million values calculated by RSEM were log2+1 transformed. A total of 401 nuclei pooled from 14 mice (9 males and 5 females) were included in the analysis.

### Single nuclei RNA-seq data preprocessing

Seurat was used for cell clustering analysis. The top 2,000 highly variable genes were determined by the variance stabilizing transformation (vst) method. Significant principal components (PCs) were estimated independently for each dataset. Clustering was performed based on these significant PCs. For UMAP plots, the default parameters were used, except the number of PCs was determined using the JackStraw() function in Seurat. Differentially expressed genes between cell types were determined using a Wilcoxon test with false discovery rate (FDR) < 0.05. GO pathway analysis was done using DAVID (https://david.ncifcrf.gov/) and significant GO terms were determined with FDR < 0.05.

### Statistics

Cell activation was analyzed in Graphpad Prism. Specific t-test or ANOVA results are described in the results section. Distributions were assumed to be normal, with no explicit normality tests performed. Significance thresholds were set at *p* < 0.05.

### Data Availability

RNA-Seq data have been submitted to GEO (accession number pending and can be made available to reviewers upon request). Other data supporting the findings of this study are available from the corresponding author upon reasonable request.

## ACKNOWLEDGMENTS

The authors thank Mary Lynn Gage for editorial assistance, Caz O’Connor and Lara Boggeman for flow cytometry guidance, and Xavier Zhou and Amanda Argoncillo for rodent technical help. This work was supported by NIH grant #MH114030 (F.H.G.), the JPB Foundation, American Heart Association/Paul G. Allen Frontiers Group Brain Health & Cognitive Impairment Initiative (19PABHI34610000), Annette C. Merle-Smith, the Dolby Family, and the Brinson Foundation. We also thank the Salk Institute Core facilities, in particular: the Waitt Advanced Biophotonics Core Facility with funding from NIH-NCI CCSG: P30 014195 and the Waitt Foundation; the Flow Cytometry Core Facility with funding from NIH-NCI CCSG: P30 014195; and the NGS Core Facility with funding from NIH-NCI CCSG: P30 014195, the Chapman Foundation and the Helmsley Charitable Trust. The authors declare no competing interests.

## FIGURE LEGENDS

**Figure S1. Full FACS gating strategy for collecting active immature and mature DGC nuclei**. Nuclei were identified based on forward and side scatter and were filtered based on pulse width to exclude doublets. GFP- (mature) and GFP+ (immature) nuclei were both gated for presence of DGC markers PROX1 and CTIP2. ARC+ cells were classified as active and ARC- as inactive. Mature GFP-cells were used to set the gate location for immature GFP+ cells, which were much less abundant.

## REFERENCES

Akers, Katherine G., Alonso Martinez-Canabal, Leonardo Restivo, Adelaide P. Yiu, Antonietta De Cristofaro, Hwa-Lin Liz Hsiang, Anne L. Wheeler, et al. “Hippocampal Neurogenesis Regulates Forgetting during Adulthood and Infancy.” Science (New York, N.Y.) 344, no. 6184 (May 9, 2014): 598–602. https://doi.org/10.1126/science.1248903.

Alberini, Cristina M., and Eric R. Kandel. “The Regulation of Transcription in Memory Consolidation.” Cold Spring Harbor Perspectives in Biology 7, no. 1 (January 1, 2015): a021741. https://doi.org/10.1101/cshperspect.a021741.

Anacker, Christoph, and René Hen. “Adult Hippocampal Neurogenesis and Cognitive Flexibility — Linking Memory and Mood.” Nature Reviews Neuroscience 18, no. 6 (June 2017): 335–46. https://doi.org/10.1038/nrn.2017.45.

Anacker, Christoph, Victor M. Luna, Gregory S. Stevens, Amira Millette, Ryan Shores, Jessica C. Jimenez, Briana Chen, and René Hen. “Hippocampal Neurogenesis Confers Stress Resilience by Inhibiting the Ventral Dentate Gyrus.” Nature 559, no. 7712 (July 2018): 98–102. https://doi.org/10.1038/s41586-018-0262-4.

Burghardt, Nesha S., Eun Hye Park, René Hen, and André A. Fenton. “Adult-Born Hippocampal Neurons Promote Cognitive Flexibility in Mice.” Hippocampus 22, no. 9 (September 2012): 1795–1808. https://doi.org/10.1002/hipo.22013.

Costa-Mattioli, Mauro, Wayne S. Sossin, Eric Klann, and Nahum Sonenberg. “Translational Control of Long-Lasting Synaptic Plasticity and Memory.” Neuron 61, no. 1 (January 15, 2009): 10–26. https://doi.org/10.1016/j.neuron.2008.10.055.

Danielson, Nathan B., Patrick Kaifosh, Jeffrey D. Zaremba, Matthew Lovett-Barron, Joseph Tsai, Christine A. Denny, Elizabeth M. Balough, et al. “Distinct Contribution of Adult-Born Hippocampal Granule Cells to Context Encoding.” Neuron 90, no. 1 (April 6, 2016): 101–12. https://doi.org/10.1016/j.neuron.2016.02.019.

Dieni, Cristina V., Angela K. Nietz, Roberto Panichi, Jacques I. Wadiche, and Linda Overstreet-Wadiche. “Distinct Determinants of Sparse Activation during Granule Cell Maturation.” Journal of Neuroscience 33, no. 49 (December 4, 2013): 19131–42. https://doi.org/10.1523/JNEUROSCI.2289-13.2013.

Dieni, Cristina V., Roberto Panichi, James B. Aimone, Chay T. Kuo, Jacques I. Wadiche, and Linda Overstreet-Wadiche. “Low Excitatory Innervation Balances High Intrinsic Excitability of Immature Dentate Neurons.” Nature Communications 7, no. 1 (April 20, 2016): 11313. https://doi.org/10.1038/ncomms11313.

Drew, Liam J., Mazen A. Kheirbek, Victor M. Luna, Christine A. Denny, Megan A. Cloidt, Melody V. Wu, Swati Jain, Helen E. Scharfman, and René Hen. “Activation of Local Inhibitory Circuits in the Dentate Gyrus by Adult-Born Neurons.” Hippocampus 26, no. 6 (2016): 763–78. https://doi.org/10.1002/hipo.22557.

Ge, Shaoyu, Chih-hao Yang, Kuei-sen Hsu, Guo-li Ming, and Hongjun Song. “A Critical Period for Enhanced Synaptic Plasticity in Newly Generated Neurons of the Adult Brain.” Neuron 54, no. 4 (May 24, 2007): 559–66. https://doi.org/10.1016/j.neuron.2007.05.002.

Ikrar, Taruna, Nannan Guo, Kaiwen He, Antoine Besnard, Sally Levinson, Alexis Hill, Hey-Kyoung Lee, Rene Hen, Xiangmin Xu, and Amar Sahay. “Adult Neurogenesis Modifies Excitability of the Dentate Gyrus.” Frontiers in Neural Circuits 7 (2013): 204. https://doi.org/10.3389/fncir.2013.00204.

Jaeger, Baptiste N., Sara B. Linker, Sarah L. Parylak, Jerika J. Barron, Iryna S. Gallina, Christian D. Saavedra, Conor Fitzpatrick, et al. “A Novel Environment-Evoked Transcriptional Signature Predicts Reactivity in Single Dentate Granule Neurons.” Nature Communications 9, no. 1 (August 6, 2018): 3084. https://doi.org/10.1038/s41467-018-05418-8.

Jessberger, Sebastian, and Gerd Kempermann. “Adult-Born Hippocampal Neurons Mature into Activity-Dependent Responsiveness.” European Journal of Neuroscience 18, no. 10 (2003): 2707–12. https://doi.org/10.1111/j.1460-9568.2003.02986.x.

Jungenitz, Tassilo, Tijana Radic, Peter Jedlicka, and Stephan W. Schwarzacher. “High-Frequency Stimulation Induces Gradual Immediate Early Gene Expression in Maturing Adult-Generated Hippocampal Granule Cells.” Cerebral Cortex (New York, N.Y.: 1991) 24, no. 7 (July 2014): 1845–57. https://doi.org/10.1093/cercor/bht035.

Kee, Nohjin, Cátia M. Teixeira, Afra H. Wang, and Paul W. Frankland. “Preferential Incorporation of Adult-Generated Granule Cells into Spatial Memory Networks in the Dentate Gyrus.” Nature Neuroscience 10, no. 3 (March 2007): 355–62. https://doi.org/10.1038/nn1847.

Kempermann, Gerd, Hongjun Song, and Fred H. Gage. “Neurogenesis in the Adult Hippocampus.” Cold Spring Harbor Perspectives in Biology 7, no. 9 (September 1, 2015): a018812. https://doi.org/10.1101/cshperspect.a018812.

Kim, Euiseok J., Jessica L. Ables, Lauren K. Dickel, Amelia J. Eisch, and Jane E. Johnson. “Ascl1 (Mash1) Defines Cells with Long-Term Neurogenic Potential in Subgranular and Subventricular Zones in Adult Mouse Brain.” PloS One 6, no. 3 (March 31, 2011): e18472. https://doi.org/10.1371/journal.pone.0018472.

Lacar, Benjamin, Sara B. Linker, Baptiste N. Jaeger, Suguna Rani Krishnaswami, Jerika J. Barron, Martijn J. E. Kelder, Sarah L. Parylak, et al. “Nuclear RNA-Seq of Single Neurons Reveals Molecular Signatures of Activation.” Nature Communications 7, no. 1 (April 19, 2016): 11022. https://doi.org/10.1038/ncomms11022.

Laplagne, Diego A., M. Soledad Espósito, Verónica C. Piatti, Nicolás A. Morgenstern, Chunmei Zhao, Henriette van Praag, Fred H. Gage, and Alejandro F. Schinder. “Functional Convergence of Neurons Generated in the Developing and Adult Hippocampus.” PLOS Biology 4, no. 12 (November 21, 2006): e409. https://doi.org/10.1371/journal.pbio.0040409.

Li, Liyi, Sébastien Sultan, Stefanie Heigele, Charlotte Schmidt-Salzmann, Nicolas Toni, and Josef Bischofberger. “Silent Synapses Generate Sparse and Orthogonal Action Potential Firing in Adult-Born Hippocampal Granule Cells.” Edited by Yukiko Goda. ELife 6 (August 8, 2017): e23612. https://doi.org/10.7554/eLife.23612.

Marín-Burgin, Antonia, Lucas A. Mongiat, M. Belén Pardi, and Alejandro F. Schinder. “Unique Processing during a Period of High Excitation/Inhibition Balance in Adult-Born Neurons.” Science (New York, N.Y.) 335, no. 6073 (March 9, 2012): 1238–42. https://doi.org/10.1126/science.1214956.

McAvoy, Kathleen M., Kimberly N. Scobie, Stefan Berger, Craig Russo, Nannan Guo, Pakanat Decharatanachart, Hugo Vega-Ramirez, et al. “Modulating Neuronal Competition Dynamics in the Dentate Gyrus to Rejuvenate Aging Memory Circuits.” Neuron 91, no. 6 (September 21, 2016): 1356–73. https://doi.org/10.1016/j.neuron.2016.08.009.

Mo, Alisa, Eran A. Mukamel, Fred P. Davis, Chongyuan Luo, Gilbert L. Henry, Serge Picard, Mark A. Urich, et al. “Epigenomic Signatures of Neuronal Diversity in the Mammalian Brain.” Neuron 86, no. 6 (June 17, 2015): 1369–84. https://doi.org/10.1016/j.neuron.2015.05.018.

Mongiat, Lucas A., M. Soledad Espósito, Gabriela Lombardi, and Alejandro F. Schinder. “Reliable Activation of Immature Neurons in the Adult Hippocampus.” PLOS ONE 4, no. 4 (April 28, 2009): e5320. https://doi.org/10.1371/journal.pone.0005320.

Picelli, Simone, Åsa K. Björklund, Omid R. Faridani, Sven Sagasser, Gösta Winberg, and Rickard Sandberg. “Smart-Seq2 for Sensitive Full-Length Transcriptome Profiling in Single Cells.” Nature Methods 10, no. 11 (November 2013): 1096–98. https://doi.org/10.1038/nmeth.2639.

Picelli, Simone, Omid R. Faridani, Asa K. Björklund, Gösta Winberg, Sven Sagasser, and Rickard Sandberg. “Full-Length RNA-Seq from Single Cells Using Smart-Seq2.” Nature Protocols 9, no. 1 (January 2014): 171–81. https://doi.org/10.1038/nprot.2014.006.

Praag, Henriette van, Alejandro F. Schinder, Brian R. Christie, Nicolas Toni, Theo D. Palmer, and Fred H. Gage. “Functional Neurogenesis in the Adult Hippocampus.” Nature 415, no. 6875 (February 2002): 1030–34. https://doi.org/10.1038/4151030a.

Rangel, L. M., A. S. Alexander, J. B. Aimone, J. Wiles, F. H. Gage, A. A. Chiba, and L. K. Quinn. “Temporally Selective Contextual Encoding in the Dentate Gyrus of the Hippocampus.” Nature Communications 5 (2014): 3181. https://doi.org/10.1038/ncomms4181.

Schmidt-Hieber, Christoph, Peter Jonas, and Josef Bischofberger. “Enhanced Synaptic Plasticity in Newly Generated Granule Cells of the Adult Hippocampus.” Nature 429, no. 6988 (May 2004): 184–87. https://doi.org/10.1038/nature02553.

Snyder, Jason S., Jessica S. Choe, Meredith A. Clifford, Sara I. Jeurling, Patrick Hurley, Ashly Brown, J. Frances Kamhi, and Heather A. Cameron. “Adult-Born Hippocampal Neurons Are More Numerous, Faster Maturing, and More Involved in Behavior in Rats than in Mice.” Journal of Neuroscience 29, no. 46 (November 18, 2009): 14484–95. https://doi.org/10.1523/JNEUROSCI.1768-09.2009.

Stone, Scellig S.D., Cátia M. Teixeira, Kirill Zaslavsky, Anne L. Wheeler, Alonso Martinez-Canabal, Afra H. Wang, Masanori Sakaguchi, Andres M. Lozano, and Paul W. Frankland. “Functional Convergence of Developmentally and Adult-Generated Granule Cells in Dentate Gyrus Circuits Supporting Hippocampus-Dependent Memory.” Hippocampus 21, no. 12 (2011): 1348–62. https://doi.org/10.1002/hipo.20845.

Tashiro, Ayumu, Hiroshi Makino, and Fred H. Gage. “Experience-Specific Functional Modification of the Dentate Gyrus through Adult Neurogenesis: A Critical Period during an Immature Stage.” Journal of Neuroscience 27, no. 12 (March 21, 2007): 3252–59. https://doi.org/10.1523/JNEUROSCI.4941-06.2007.

Toda, Tomohisa, Sarah L. Parylak, Sara B. Linker, and Fred H. Gage. “The Role of Adult Hippocampal Neurogenesis in Brain Health and Disease.” Molecular Psychiatry 24, no. 1 (January 2019): 67–87. https://doi.org/10.1038/s41380-018-0036-2.

Toni, Nicolas, Diego A. Laplagne, Chunmei Zhao, Gabriela Lombardi, Charles E. Ribak, Fred H. Gage, and Alejandro F. Schinder. “Neurons Born in the Adult Dentate Gyrus Form Functional Synapses with Target Cells.” Nature Neuroscience 11, no. 8 (August 2008): 901–7. https://doi.org/10.1038/nn.2156.

Wu, Ye Emily, Lin Pan, Yanning Zuo, Xinmin Li, and Weizhe Hong. “Detecting Activated Cell Populations Using Single-Cell RNA-Seq.” Neuron 96, no. 2 (October 11, 2017): 313-329.e6. https://doi.org/10.1016/j.neuron.2017.09.026.

Yang, Sung M., Diego D. Alvarez, and Alejandro F. Schinder. “Reliable Genetic Labeling of Adult-Born Dentate Granule Cells Using Ascl1CreERT2 and GlastCreERT2 Murine Lines.” Journal of Neuroscience 35, no. 46 (November 18, 2015): 15379–90. https://doi.org/10.1523/JNEUROSCI.2345-15.2015.

Yap, Ee-Lynn, and Michael E. Greenberg. “Activity-Regulated Transcription: Bridging the Gap between Neural Activity and Behavior.” Neuron 100, no. 2 (October 24, 2018): 330–48. https://doi.org/10.1016/j.neuron.2018.10.013.

Zhao, Chunmei, E. Matthew Teng, Robert G. Summers, Guo-li Ming, and Fred H. Gage. “Distinct Morphological Stages of Dentate Granule Neuron Maturation in the Adult Mouse Hippocampus.” Journal of Neuroscience 26, no. 1 (January 4, 2006): 3–11. https://doi.org/10.1523/JNEUROSCI.3648-05.2006.

