## Supplementary figures and images for "Neuronal activity-related transcription is blunted in immature compared to mature dentate granule cells"

### Supplemental Figure 1

**Figure S1: Full FACS gating strategy for collecting active immature and mature DGCs**

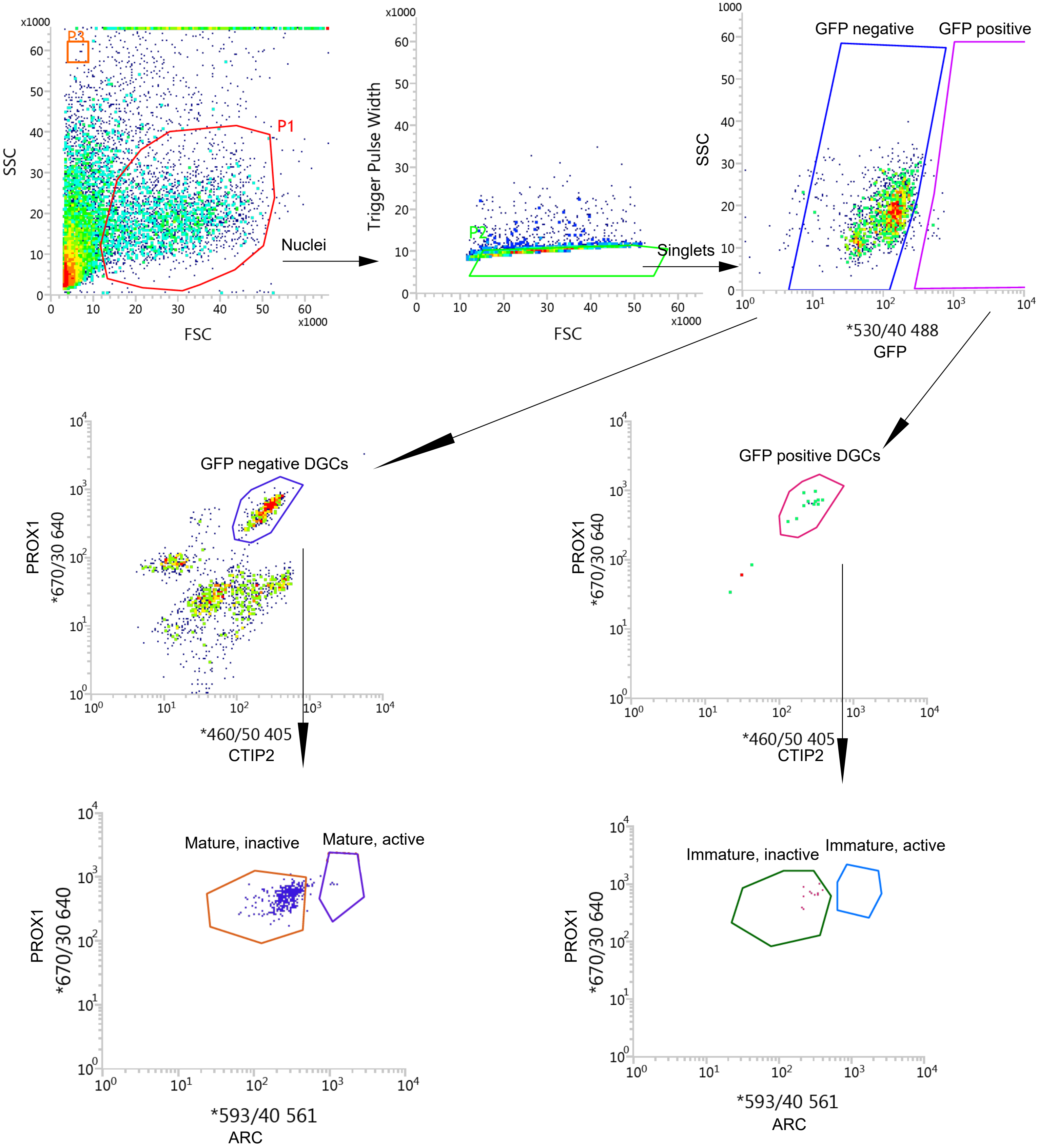
